# Tiny radio-tag backpacks impact, but do not significantly affect, hummingbird time budgets in captivity

**DOI:** 10.1101/2025.01.29.635563

**Authors:** Alyssa J. Sargent, Ana Melisa Fernandes, Aeris E. Clarkson, Samantha-Lynn Martinez, Alexandra Coenen, Laney Hansell, Yash P. Talwekar, Miguel A. Muñoz-Amaya, Nicolas Téllez-Colmenares, Rosalee Elting, Yutong Sun, Olivia A. Cartwright, Nicole Büttner, Alejandro Rico-Guevara

## Abstract

**Background:** Given that wildlife tags have recently become miniaturized enough to work with some of the world’s smallest vertebrates, there is a newfound urgency for affordable, field-accessible biologging ethics studies. We designed a 3-hour time-budget experiment to investigate how radio-transmitter backpacks affect hummingbirds’ behavior. Using a large flight arena in Colombia, we individually filmed 25 Black-throated Mangoes (*Anthracothorax nigricollis*) under two randomized treatments, tagged and untagged, to characterize and quantitatively compare their behavior. We analyzed all videos using the Behavioral Observation Research Interactive Software (BORIS), to create time-budget breakdowns of our key behaviors of interest: flying, feeding, preening, and perching. We also designed an aviary-style “entanglement test” (n = 30) to determine if any individuals would snag on vegetation while equipped with the backpack harness, and tested 6 additional birds in this enclosure overnight for any longer-term negative effects.

**Results:** Across duration, number of bouts, and bout length, we found no significant differences in the behavior of individuals (flying, feeding, preening, and perching) when they were or were not tagged. However, the additive effects of treatment number (whether the bird was undergoing its first or second 1.5-hour treatment) and treatment type (tagged or untagged) most accurately predicted time spent flying (birds flew significantly more in their second 1.5-hour treatment). The weight of the bird, meanwhile, best predicted feeding duration (lighter birds fed significantly more). Lastly, the additive effects of time of day and treatment type had the highest predictive accuracy of time spent preening (birds preened significantly more in the afternoon than the morning, and significantly more in the evening than the afternoon); here, the effect of treatment type was highly significant. In our aviary tests, no individuals became entangled in vegetation or exhibited any adverse overnight effects from harness wear.

**Conclusions:** In our captive study, radio-transmitter backpacks did not significantly affect hummingbird behavior when considered independently; however, additional covariates are essential to account for, and the effects of being in a confined space may also be significant. Nonetheless, our experimental model is relatively straightforward to fine-tune to other small taxa and is suitable for remote conditions, providing a useful basis with which to examine species-specific effects of biologging prior to starting field studies.

## INTRODUCTION

As biologging devices become ever-smaller and increasingly prevalent, the number of studies on animal movement, behavior, and physiology has risen substantially, providing a wealth of environmental and ecological information [1,2]. This surge has brought **biologging ethics**—the study of animals’ welfare while equipped with auxiliary devices—to the forefront of automated tracking concerns, and for good reason [3–6]. Biologgers have been tied to a slew of negative effects, impacting behavior [7], sensory organs [8], foraging [9], reproduction [9–11], energetics [9,12], and survival [11,13,14].

Device weight is commonly considered the root of many of these issues and, as a means of addressing them, many biologging studies conform to the “3–5% rule”: a widely accepted ethical standard wherein devices must weigh below 3–5% of the animal’s bodyweight [15–17]. Many researchers have, however, criticized this rule for its lack of evidential support [6,10,14,18,19] and its overly simplistic approach [19]. *How* and *where* a biologger is attached to an animal can also present complications [13,14,20]—and the 3–5% rule accounts for neither biomechanics (e.g., relative wing area to body size) nor life history (e.g., aquatic vs. terrestrial lifestyles [3,14]). Volant animals, in particular, must compensate for the additional weight and drag of tracking devices by adjusting their flight kinematics (e.g., increasing stroke amplitude and/or wingbeat frequency [21,22]), which not only increases energy expenditure, but also reduces performance [6,10].

Nevertheless, many avian biologging studies do not report or test for tag effects [4]; there is still a pressing need for species-specific experimental designs [23,24]. Though tools such as respirometers and wind tunnels can provide highly detailed information on device effects, they do not provide any information regarding birds’ potential for entanglement in vegetation or longer-term impacts (e.g., multi-hour assessments across which many more behaviors can be evaluated); furthermore, this equipment can be both costly and complex [25,26], and is not always suitable for the remote field settings where many automated tracking studies take place [2]. Conversely, **time budgets** (which break down the proportion of time an animal spends conducting various behaviors) can act as a useful, field-accessible tool for biologging welfare studies. As a biologger may increase the energy needed to perform a specific activity, or prevent it entirely [3], time budgets allow researchers to compare between tagged and untagged individuals from the same population [4,5,24].

Hummingbirds are notoriously energy-constrained birds [27–29]; as such, researchers applying auxiliary devices to hummingbirds should consider welfare and time budgeting of critical importance (see [24] for a fundamental example). It is well-established that hummingbirds are capable of remarkable feats of strength: lifting well over their bodyweight in maximum performance tests (e.g., [30,31]); sustaining flight in heavy rain, a mere 20 seconds of which can increase their bodyweight by 20% [32]; and, in the case of females, developing and carrying eggs up to 15% of their own bodyweight [33]. However, these displays of burst or short-term power may not be indicative of more prolonged weight-carrying capabilities, and/or the ability to fly with external aerodynamic impediments [2]. Biologgers have only recently miniaturized enough to tag these tiny birds [24,34–37], and ethical studies of their effects have been sparse [24]. The fresh introduction of backpack harnesses to hummingbird research [35] has allowed for an advantageous combination: tags with lengthy detection distances (unlike passive integrated transponders [38,39]) that can be deployed for far longer periods than those affixed with glue (such as battery-powered radio-transmitters [34]). Yet hummingbird backpacks are still relatively novel, and we lack any ethical quantifications of how they affect the behavior of individual birds.

With the hypothesis that wearing tags is costly for hummingbirds [24], we designed a time-budget experiment to determine the behavioral impacts of a backpack harness with a custom chassis, along with a field-based test to examine hummingbirds’ propensity to get entangled in vegetation while fitted with it (see [40] for a documented case of entanglement and mortality in Hawaiian honeycreepers when equipped with radio-transmitters). In this study, we compared the behavior of hummingbirds when they were equipped with a radio-transmitter versus when they were not. We predicted that, when tagged, individuals would fly less to minimize energetic expenditure [24]—and, by extension, perch more—as well as feed more to maximize energetic intake from *ad libitum* syringes (but see [41] for the opposite results in a free-living study of goldeneyes), and preen more in attempts to physically remove the radio-transmitters [41,42].

## METHODS

### Study Overview

After conducting a pilot-iteration of this study on the Green-crowned Brilliant (*Heliodoxa jacula*), White-necked Jacobin (*Florisuga mellivora*), and White-tipped Sicklebill (*Eutoxeres aquila*) in Ecuador (detailed in **Additional File 1: Ecuador Pilot**, **Table S1**), we conducted all of our experiments in the dry season of August and September 2022 at Centro de Investigación Colibrí Gorriazul (hereafter referred to as “Gorriazul”) in Colombia. Gorriazul is situated at roughly 1,700 m elevation, midway up the western slope of the Eastern Cordillera of the Colombian Andes, in a matrix of pristine rainforest and agricultural properties. At Gorriazul, there is an established network of ∼20 sugar-water feeders and a long-standing, diverse population of hummingbirds of roughly 12–18 species, which feed from both these feeders and many native flowers in the surrounding landscape.

We chose to work with the Black-throated Mango (*Anthracothorax nigricollis*), on account of its large body size (7–9 g) and significant abundance onsite; to further reduce confounding effects that may impact behavior, we worked selectively with adult birds that had typically male-associated plumage. It is worth noting that there are recorded cases of female polymorphism— “male-like females”— within this species [43–45], but we lacked the resources to genetically sex individuals. We captured all birds using baited drop-net traps [46], either processing them immediately post-capture or rapidly placing them in bird bags as a form of sensory deprivation to reduce stress [3].

### Tagging Procedure

We first identified, aged (based on plumage and bill corrugations), sexed, took baseline morphometric measurements for, and scored the body conditions of each newly captured bird [46]. See **Table 1** for all bodyweight and tag specifications. Next, we injected each individual with an interscapular, subcutaneous passive integrated transponder (“PIT tag”: BioMark) and closed the insertion site with surgical glue (VetBond) as described by Bandivadekar et al. [38]. These 8 mm, 35 mg PIT tags proved to be more informative identifiers than tarsal bands—for which hummingbird-specific tools are prohibitively difficult to access or fabricate, especially given the lack of methods standardization for tropical species. After implantation, we released each bird and monitored its return to our feeders, which were equipped with radio-frequency identification (RFID) antennas. By cross-checking RFID readings of PIT tags with capture data, we selectively recaptured individuals after a minimum of 7 days, weighed them, and verified that insertion wounds had healed before proceeding with further experimentation.

**Table 1.** Black-throated Mango bodyweights and tag specifications, with means and standard deviations, across all trials (enclosure and aviary).

| Item | Mean weight (grams) | Mean Percent bodyweight |
| --- | --- | --- |
| Individual before trial | 7.92 ± 0.55 | - |
| Individual after trial | 8.26 ± 0.64 | - |
| Individual across trial (averaged) | 8.10 ± 0.53 | - |
| CTT tag | 0.39 ± 0.01 | 4.87 ± 0.30 |
| CTT tag + harness | 0.44 ± 0.02 | 5.49 ± 0.40 |
| CTT tag + harness + PIT | 0.48 ± 0.02 | 5.93 ± 0.42 |

We then equipped each individual with a radio-transmitter (Cellular Tracking Technologies “LifeTag”) backpack as per the experiments described below, replicating the process of Williamson and Witt [35] and spending approximately 5–7 minutes per harness application (we restrained the birds using a strip of cloth and a small clip for the final double surgeon’s knot, **Additional File 1: Fig. S1**). When prepared with our custom chassis and attachment materials (**Fig. 1**), these radio-transmitters gained, on average, 53 mg in weight. See **Additional File 1** (**Harness Design**) for an in-depth description of our chassis fabrication process, which we optimized for ultra-lightweight and durable field applications, as well as a discussion of its practical benefits.

**Figure 1.**
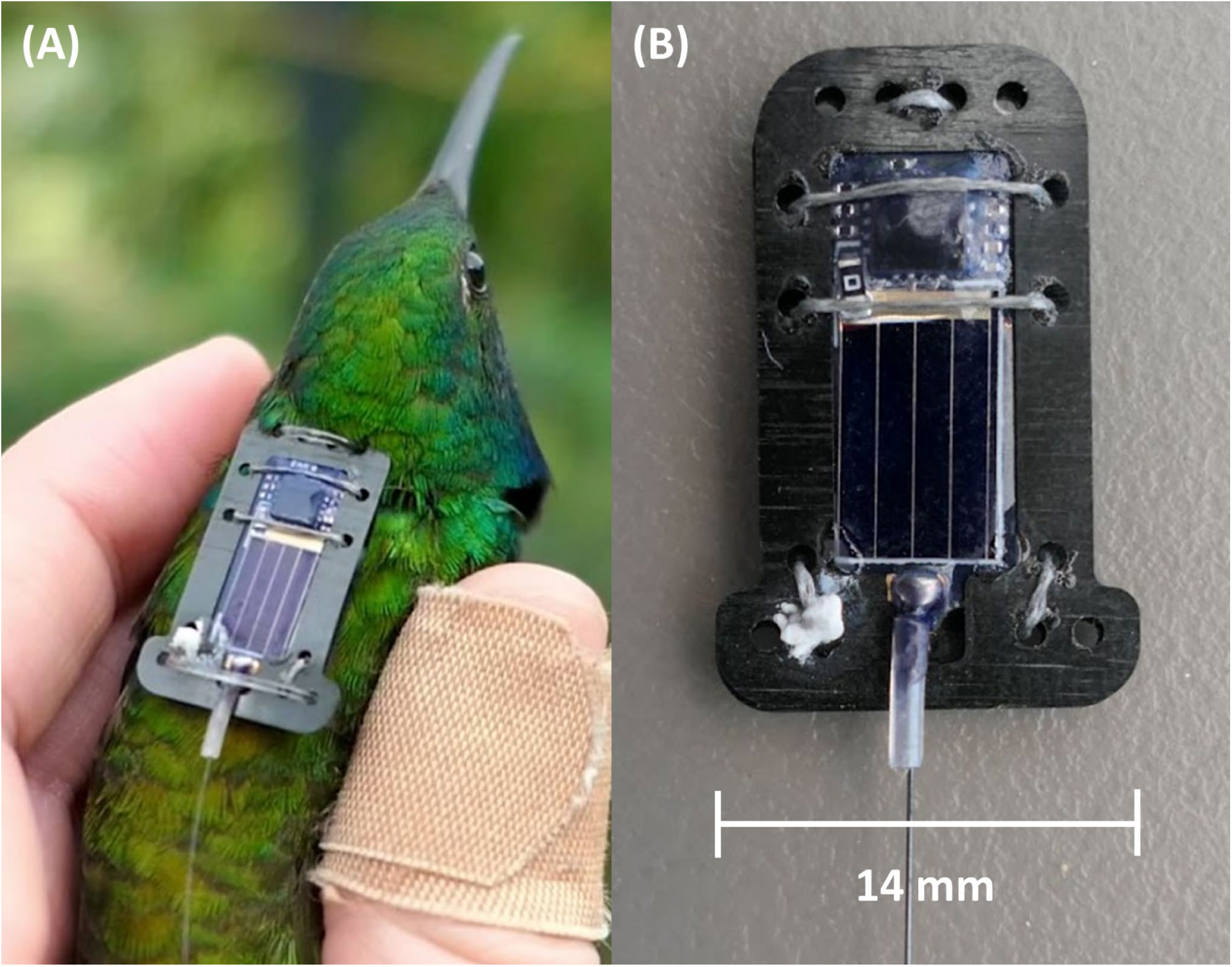
**(A**) Adult male Black-throated Mango wearing radio-transmitter backpack with custom chassis. **(B)** Close-up view of completed chassis attachment (composed of carbon fiber reinforced polymer) on solar-powered radio-transmitter. Tag ID is cut out of the bottom chassis component; solar cell is left exposed. Four corner holes left open for the StretchMagic harness and white mark (bottom left) is dried superglue used to secure the fishing line knot.

### Enclosure Experimental Design

We designed a 3-hour, “enclosed-environment” time-budget experiment to monitor hummingbird weight, feeding, and behavior in a controlled space. To establish baseline behavior in captivity and quantify the “degree of departure from the norm” [3] when tagged, we tested each individual (n = 25) under two treatments, randomized in order: tagged and untagged with a radio-transmitter harness (following similar methodologies established by Zenzal et al. [24]). After offering each individual nectar from a handheld syringe to prompt its flower-like recognition, we began the experimental procedure: for each 1.5 hour treatment, we released the bird into a flight tent (setup detailed below), allowed it 30 minutes to acclimate, and then filmed its behavior for 1 hour. We defined *acclimation failure* as a bird displaying signs of distress (remaining stationary on tent floor or gaping bill, as characterized by Russell et al. [46]), and/or an inability to identify either the nectar-filled syringe or the perch we provided, in these first 30 minutes. In such cases, we terminated the trial early, then hand-fed and released the individual. If there was adequate time after completing a full trial (the term *trial* refers to the entire 3-hour experimental procedure for a single individual), we subsequently tested the bird in a one-hour aviary test (**Methods: Aviary Tests**), completing all work at least 1 hour prior to local sunset to provide each individual a sufficient period to feed and readjust in the wild before roosting. In total, 22 of our birds underwent a 3-hour trial and a subsequent aviary test.

Our flight arena was a six-person Coleman tent (3.0 x 2.4 x 1.8 m) that we pitched directly inside the research center. Inside, we placed a tripod mounted with two Raspberry Pi load cells, one for a feeder syringe (provided sucrose solution was 15–20% w/w, measured with a refractometer at the beginning of each trial) and one for the only perch available (**Additional File 1: Fig. S2**). We also equipped the tent with **a)** a GoPro Hero 7 Black (720p resolution) in the side pocket to surveil each bird via live feed, to ensure that it fed and perched successfully without displaying signs of distress (see above); and **b)** a thermometer with a wire sensor threaded through the door, to monitor interior temperature and ensure that it did not get too hot throughout the day (our temperatures did not exceed 28.5 °C, remaining below the 32 °C maximum recommended by Russell et al. [46]). We separated the tent from researchers via a dividing sheet (draped from the ceiling) in the middle of the room, to prevent visual stimulation and further stress for the birds [3]. We used a Minolta MN80NV camcorder (1080p resolution, 30 fps) and a backup JVC camera (1080p resolution, 60 fps), which performed well without overheating during our extended filming periods in tropical conditions. Additionally, owing to substantial drift from the load cells, we elected to weigh each individual and the syringe before and after each trial instead; we also checked all individuals for any sign of harness-induced skin abrasion at the conclusion of each trial.

### Enclosure Analysis

We analyzed all treatment videos frame-by-frame using the Behavioral Observation Research Interactive Software [47], which creates time-budget breakdowns from logged behaviors of interest. Through BORIS, we calculated the total number of seconds individuals spent performing specific behaviors—flying, hover-feeding, preening, and perching (**Additional File 2: Video S1**)—over the course of each given treatment’s (tagged or untagged) observation period. As we began video-recording immediately after each 30-minute acclimation period, regardless of what the bird was doing, frame-counting could have begun while the bird was conducting any behavior (e.g., mid-flight). We defined the start and end of each ethological category as the first and last frames, respectively, in which the bird visibly changed its behavior. As such, we considered the bird to be “hover-feeding” whenever its bill was in contact with the syringe feeder, “preening” at any time that its bill was in contact with its feathers (following Zenzal et al. [24]), “perching” at any point when its feet were in contact with the perch, and “flying” whenever the bird was airborne but not hover-feeding. We conducted all remaining analyses using R version 4.3.1 [48].

### Enclosure Tag Effects

We first assessed whether the durations of each behavior were distributed parametrically using the Shapiro-Wilk normality test. To test our predictions directly, we conducted one-tailed, paired-sample tests against our alternative hypotheses (see **Introduction**): for parametric distributions, t-tests (*stats* package [48]), and for non-parametric, either asymptotic Wilcoxon-Mann-Whitney tests (*coin* package [49]) or Wilcoxon signed-rank tests (*stats* package [48]) given the presence or absence of ties, respectively. In these tests, we separately compared the time and percent of time that individuals spent performing each key behavior—flying, hover-feeding, preening, and perching without preening (perching duration minus preening duration, to result in percentages that added up to 100)—when tagged and untagged. We also repeated this analytical process with the number of behavioral “bouts” that individuals performed for each treatment, along with the average length of these bouts; here, we did not adjust perching values to account for when the bird was or was not preening, so as to quantitatively characterize “full” perching bouts.

### Enclosure Model Selection

Using the *lme4* [50], *lmerTest* [51], and *glmmTMB* [52] packages, we fit a series of general and generalized linear mixed-effects models to test whether radio-transmitters, or any other independent variables (see below), would best explain the *durations* of the behaviors that we observed. As the combined durations of flying and hover-feeding were the inverse of perching duration within each 3,600-second observation period—the behaviors are mutually exclusive—we did not include time spent perching in our regression analyses, and rather focused on flying, hover-feeding, and preening durations as our dependent variables. Incorporating a random effect of individual each time, we individually compared each of the following fixed effects in a separate model: **treatment type** (2 levels: tagged and untagged); individual’s **weight** at the start of the trial (continuous, in grams); **treatment number** (2 levels: treatment 1 and treatment 2); tent’s average internal **temperature** for the treatment (continuous, in Celsius); **percent sucrose** of the provided nectar throughout the trial (continuous); and **period of day** wherein the majority of the treatment took place (3 levels: morning, 6:00–11:59; afternoon, 12:00–15:59; and early evening, 16:00–on). For each fixed effect that performed better than the null (see below), we also ran an additive and interactive model that incorporated treatment type alongside it (e.g., treatment type + period of day, treatment type * period of day).

Using the *DHARMa* package [53], we checked for the most appropriate model error structure for each behavioral category through dispersion, outlier, homoscedasticity, and zero-inflation tests; we also assessed the structure and distribution of our residuals with QQ-normality and residual vs. fitted-value plots. While a Gaussian distribution provided the optimal fit for flight models, feeding and preening models were best suited for a zero-inflated tweedie distribution— although the tweedie model’s performance worsened when it came to predicting longer preening durations, given the extreme level (44%) of zero inflation (frequent zero counts).

We conducted all model selection using Akaike’s information criterion corrected for small sample sizes, seeking models that performed better (ΔAICc < 2) than the null—which only included the random effect of individual—or better than the model containing our primary fixed effect of interest, treatment type. We also assessed the significance of top model terms with likelihood ratio tests using the *lmtest* package [54]. For top models, we present the p-values and effect sizes (β-estimates) of individual parameters; if their 95% confidence intervals did not overlap zero, we considered them to be significant influences on behavior [55].

### Enclosure Inter-Rater (Observer) Reliability

To determine the reliability of our ethological video analyses and the protocols with which we differentiated behavior types, we used the *irr* package to conduct an inter-rater reliability analysis [56], which is used to quantify the level of consistency between different raters (observers). As per Harvey [57], continuous data call for an intra-class correlation coefficient; we calculated this statistic using a consistency type, single-measure, two-way random-effects model for 24 measurements (6 raters scoring the same 4 videos). Here, we compared raters’ calculated durations (of flying, hover-feeding, preening, and perching) for the subset of videos in a single statistical test.

### Aviary Tests

We designed two “open-environment” aviary tests—a 1-hour test (n = 30, 22 of which underwent a prior enclosure experiment) and an overnight test (n = 6)—to monitor the flight of harnessed hummingbirds while exposed to natural vegetation. Our aviary consisted of a large, open-floored, mesh-walled tent (Tailgaterz, 3.3 x 2.7 x 2.0 m), pitched outside (**Additional File 1: Fig. S3**) and containing 5 hummingbird feeders in syringe- and bottle-form. To identify any harness-induced entanglement [40], we also ensured that the tent contained numerous shrubs and large branches, which doubled as both perches and flight obstacles.

In the case of the 1-hour trials, we monitored individuals continuously to verify active flight within the tent without ensnarement; when conducting overnight trials, we ensured that all birds were acclimated to the tent at least one hour prior to sunset, then monitored individuals until they successfully fed from the feeders, and then once every hour until sunset. Once we had determined that the bird had fallen asleep, we left it undisturbed until early morning, at which point we checked the bird for any signs of harness-induced abrasion or chafing, removed the backpack harness, weighed the individual, and released it—except in the case of a single, extended trial that we ran for 24 hours, wherein we resumed hourly checks until an early afternoon release. In all cases, we monitored individuals from afar (with the occasional aid of binoculars) to reduce their stress levels, and treated our recapture efforts at the end of each trial as a proxy for tagged individuals’ ability to evade predators without entanglement. As these trials were purely observational with a binary outcome (successful free flight or ensnarement), we did not conduct a statistical analysis on this component of the study.

## RESULTS

### Time-Budget Summaries and Tag Effects

We had two cases of acclimation failure, in which one bird failed to feed from the syringe and a second failed to locate and rest upon the perch; subsequently, both individuals displayed signs of distress in the first 30 minutes of the trial. As such, we did not include either bird in our dataset (although we note that neither suffered lasting distress, and flew away vigorously upon their immediate release). With 25 successful trials each containing two 1-hour video-recorded observation periods, our complete dataset comprised 50 total hummingbird observation hours. Over the course of the 3-hour trials, these individuals drank an average of 3.65 ± 1.62 grams of sugar water from the syringe—nearly half their bodyweight. From start to finish, the birds themselves experienced weight changes ranging from a loss of 1.13 grams to a gain of 1.82 grams; their average change in weight, however, was an increase of 0.48 grams (standard deviation ± 0.59 grams). In no cases did the hummingbirds’ flight patterns, standard forward flight and hovering alike, appear impeded by the addition of the radio-transmitter backpack harness.

While feeding durations proved to be normally distributed (Shapiro-Wilk p = 0.354), flight, preening, and perching without preening were not (Shapiro-Wilk p = 3.662e-05, 1.808e-10, and 3.95e-04, respectively). The proportion of the treatment period that individuals spent conducting each behavior differed when the bird was tagged or not (**Table 2**, **Fig. 2**): when tagged, individuals spent on average 3.46% less time flying, 0.19% more time hover-feeding, 1.48% more time preening, and 1.78% more time perching without preening. However, none of these differences were significant as per our paired-sample t-tests, Wilcoxon signed-rank tests, or asymptotic Wilcoxon-Mann-Whitney tests. We present these results in **Table 2**.

**Figure 2.**
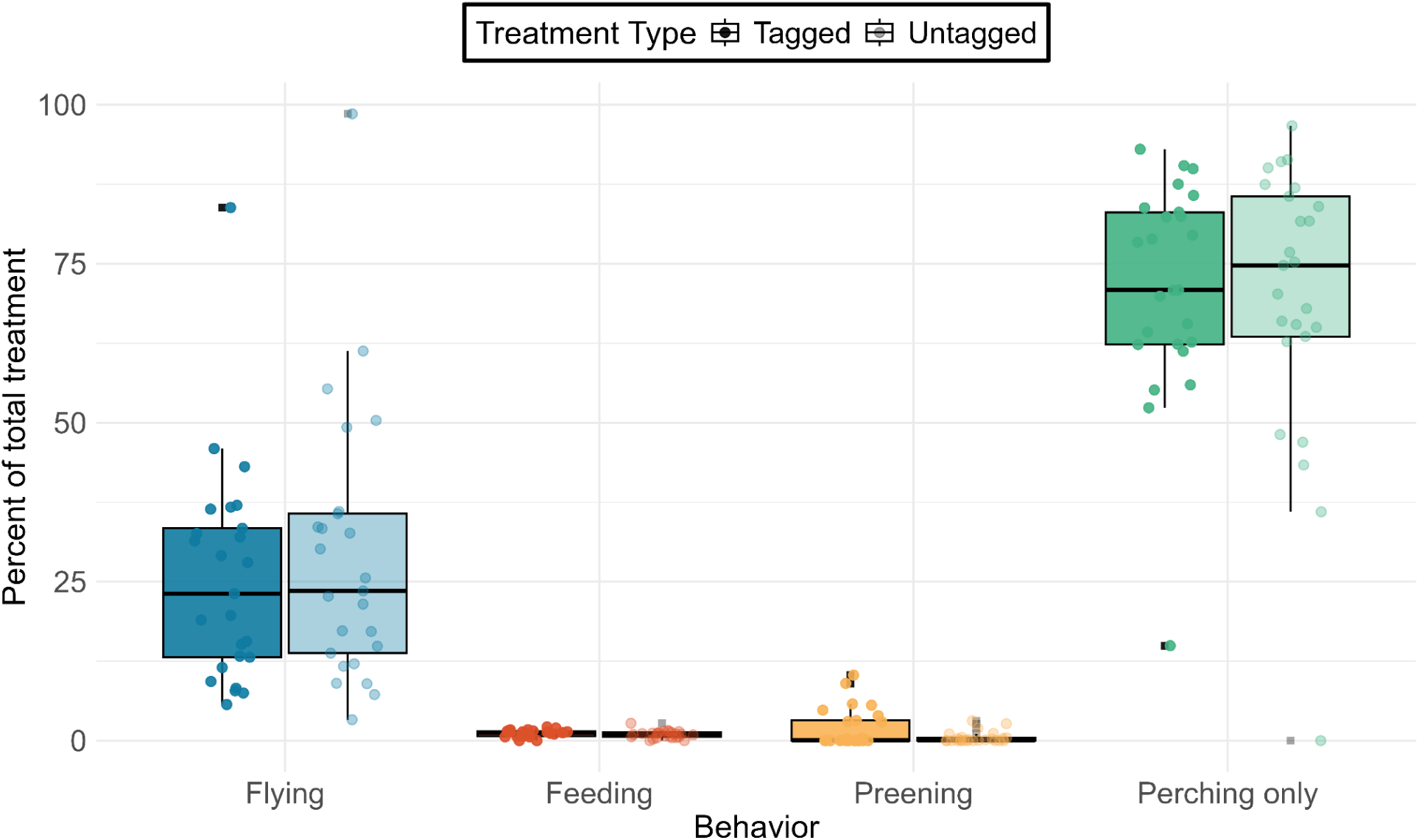
Average percentage of time that individuals spent performing various behaviors during their tagged (darker colors, to the left-hand side for each pair) and untagged (lighter colors, right-hand side) treatments. Note that “perching only” refers to perching without simultaneous preening.

**Table 2.** Total durations and percent of time (mean and standard deviations) that individuals spent conducting key behaviors during each treatment (tagged and untagged), as well as one-tailed, paired-sample test (t-test, Wilcoxon signed-rank, and asymptotic Wilcoxon-Mann-Whitney) results for each.

|  | Tagged |  | Untagged |  |  |  |  |
| --- | --- | --- | --- | --- | --- | --- | --- |
| Behavior (Wilcoxon) | Duration (s) | Percentage | Duration (s) | Percentage | V | p |  |
| Flying | 890.34 ± 612.27 | 25.54 ± 17.19 | 1030.98 ± 748.74 | 29.00 ± 21.37 | 214 | 0.0865 |  |
| Perching Without Preening | 2499.54 ± 634.62 | 71.33 ± 16.87 | 2485.74 ± 782.04 | 69.55 ± 21.75 | 155 | 0.4266 |  |
| Behavior (Mann-Whitney) | Duration (s) | Percentage | Duration (s) | Percentage | Z | p |  |
| Preening | 69.33 ± 108.07 | 1.97 ± 3.03 | 17.31 ± 30.48 | 0.49 ± 0.86 | 0.598 | 0.7252 |  |
| Behavior (paired t) | Duration (s) | Percentage | Duration (s) | Percentage | df | t | p |
| Hover-Feeding | 40.12 ± 19.42 | 1.15 ± 0.55 | 34.19 ± 20.39 | 0.96 ± 0.58 | 24 | -1.207 | 0.1196 |

In **Tables 3** and **4**, we present the average number of times individuals flew, fed, preened, and perched in each treatment (number of “bouts”), and the lengths of those bouts, respectively. In each table, we include the results from Wilcoxon signed-rank tests and asymptotic Wilcoxon-Mann-Whitney tests, as no bout distribution was parametrically distributed (all p-values from Shapiro-Wilks tests < 0.0335). In no case was the difference between bouts (number or duration) significant between tagged and untagged individuals.

**Table 3.** Number of times that individuals conducted key behaviors (“bouts”) when tagged and untagged (mean and standard deviation for both), compared with one-tailed Wilcoxon signed-rank or asymptotic Wilcoxon-Mann-Whitney tests.

| Behavior (Wilcoxon) | Number of Tagged Bouts | Number of Untagged Bouts | V | p |
| --- | --- | --- | --- | --- |
| Flying | 339 ± 230 | 362 ± 241 | 173 | 0.3957 |
| Perching | 322 ± 231 | 346 ± 231 | 181 | 0.6925 |
| Behavior (Mann-Whitney) | Number of Tagged Bouts | Number of Untagged Bouts | Z | p |
| Hover-Feeding | 18 ± 14 | 19 ± 20 | -0.117 | 0.4536 |
| Preening | 22 ± 41 | 10 ± 16 | 0.274 | 0.6079 |

**Table 4.** Length, in seconds, for each behavioral “bout” conducted by individuals when tagged and untagged (mean and standard deviation for both), compared with one-tailed Wilcoxon signed-rank or asymptotic Wilcoxon-Mann-Whitney tests.

| Behavior (Wilcoxon) | Duration of Tagged Bouts (s) | Duration of Untagged Bouts (s) | V | p |
| --- | --- | --- | --- | --- |
| Flying | 8.44 ± 23.18 | 9.01 ± 30.87 | 219 | 0.0668 |
| Perching | 284.88 ± 794.19 | 16.68 ± 25.92 | 123 | 0.1498 |
| Behavior (Mann-Whitney) | Duration of Tagged Bouts (s) | Duration of Untagged Bouts (s) | Z | p |
| Hover-Feeding | 2.88 ± 2.52 | 2.34 ± 1.46 | 0.752 | 0.6793 |
| Preening | 2.81 ± 4.77 | 1.12 ± 1.53 | 0.314 | 0.6234 |

### Model Selection

We present all our model-ranking results, each of which accounted for the random effect of individual, in **Additional File 1: Table S2**, and the parameter summaries of our top-performing models in **Additional File 1: Table S3**. Of all covariates, the amount of time that birds spent flying was most accurately predicted by the additive effects of treatment number and treatment type. On average, individuals spent 30.59% of their second treatment flying, as opposed to 23.95% in their first—a significant increase of 6.64% (p = 0.009; 95% CI = 62.64 – 374.84; **Fig. 3A**). Nonetheless, the effect of whether or not the bird was tagged was non-significant (p = 0.062; 95% CI = −6.72 – 305.49). In the case of feeding duration, bird weight had the highest predictive accuracy. In general, lighter birds fed significantly more (p = 0.009; 95% CI = −0.51 – −0.07; **Fig. 3B**). Preening durations, meanwhile, were most accurately predicted by the time of day with the additive effect of treatment type. As the day went on, the birds spent progressively more time preening: on average, birds spent 0.24% of their treatments preening when they occurred in the morning, 1.19% when they were in the afternoon, and 3.81% in the evening (**Fig. 3C**). This constituted a significant, 0.96% increase from morning to afternoon (p = 0.001; 95% CI = −2.94 – −0.74), and a 2.62% (non-significant) increase from afternoon to evening (p = 0.075; 95% CI = −0.08 – 1.72). In this model, the effect of treatment type was highly significant (p = 0.0002, 95% CI = −2.36 – −0.72).

**Figure 3.**
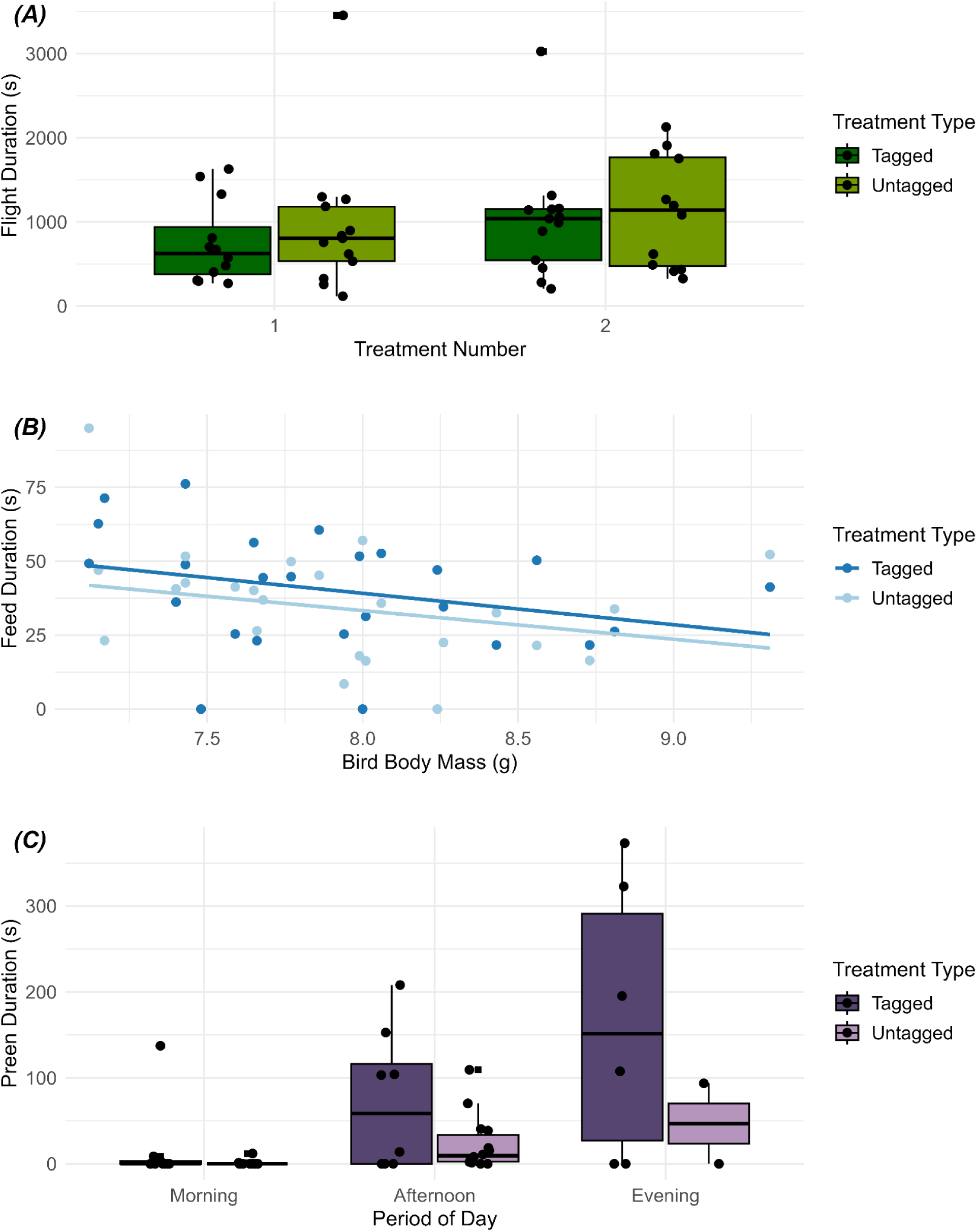
Plots showing differences in time spent performing specific behaviors when tagged and untagged relative to the fixed effect in the top model. **(A)** Flying and treatment number. **(B)** Feeding and bird body mass. **(C)** Preening and period of day. Note that each plot’s y-axis has a unique range based on the behavior in question.

For all three behavioral classes, the importance of these fixed effects was reinforced by significant likelihood ratio tests (“Flight ∼ Treatment Number + Treatment Type” p-value = 0.009, “Feeding ∼ Bird Weight” p-value = 0.001, “Preening Time of Day + Treatment Type” p-value = 3.029e-05); see **Additional File 1: Table S2**. Only in the case of flight behavior did another model (Flight ∼ Treatment Number) exhibit a competitive standing within 2 ΔAICc of our top models. In no instance was treatment type alone (i.e., not additive or interactive with another fixed effect) the top model, or competitive against the one that was.

### Inter-Rater Reliability

With 6 raters scoring the same 4 videos, our single-score intraclass correlation coefficient was 0.987 (p = 3.84e-53, 95% CI = 0.973 – 0.995), exceeding the 0.80 threshold specified by Gwet [58] and fulfilling “excellent” consistency standards [57].

### Aviary Tests

In all aviary tests, the hummingbirds flew, perched, fed, preened, and slept without getting entangled in vegetation, including during our recapture efforts, nor did they show any signs of distress [46]. Our overnight trials lasted from 15.4–24 hours (average of 18.57 ± 3.06 hours). Similarly to the enclosure experiments, there were no perceptible tag effects; the birds appeared to fly without issue. With a more spacious, natural environment, the hummingbirds were less constrained and could perform longer forward flights, as well as explore their surroundings to a greater degree. To this end, the birds appeared visibly calmer, perching and feeding more readily than their enclosure-experiment counterparts. We had no cases of acclimation failure in these tests, nor did we observe any signs of harness-induced chafing or abrasions.

## DISCUSSION

### Enclosure Experiments

As the development of biologgers (e.g., accelerometers, radio-telemeters, temperature loggers, GPS units) continues to progress at a rapid pace, the number of wildlife tracking studies will rise in turn [1,17]; we now have the technological capacity to study increasingly smaller study species in increasingly informative ways [5]. Indeed, in the three years since these experiments were conducted, commercially available biologgers have advanced significantly. Quantifying the myriad effects of devices, both in terms of attachment techniques and their presence on animals, must also continue to be a research priority in the coming years [3]. Here, our work provides a straightforward experimental design for field-accessible biologging ethics, which can easily be reproduced with many taxa: randomized, tagged and untagged treatments that incorporate both acclimation and video-recording periods for subsequent time-budget breakdowns.

To our knowledge, this is the first investigation to combine the use of multiple biologgers (PIT tags and radio-transmitters) to the study of hummingbirds, and possibly volant animals at large (see [59] for similar tests with a dummy bat). The collective weight of these two tags was, on average, 5.93% of our tested Black-throated Mangoes’ bodyweights (**Table 1**). While exceeding the 5% guideline [60], we note that hummingbirds, on account of their small size, have a higher strength-to-bodyweight ratio than larger birds and are therefore proportionally stronger [61–63]. Interestingly, while individuals did on average behave according to our predictions when equipped with radio-tag backpack harnesses (flying less, but hover-feeding, preening, and perching more), none of these differences were statistically significant—a finding that was consistent across total duration, bout number, and bout duration of each major behavior. Although Zenzal et al. [24] found that male Ruby-throated Hummingbirds spend less time flying when outfitted with a variety of radio-tags (a decrease of up to 11%, far higher than the observed 3.46% decrease in our experiment), such differences were only statistically significant in the case of heavier tags as opposed to lighter ones (6.32% and 5.79% bodyweight, respectively), the latter of which were much more similar to the proportional weight of our own.

In our experiment, accounting for other variables—either with or without the additive effect of radio-transmitter backpacks—best explained the hummingbirds’ behavior. Time spent flying, for instance, was most accurately predicted by the additive effects of treatment number (whether the bird was undergoing its first or second treatment) and treatment type (tagged or untagged). In other words, if we compared the time budgets of tagged and untagged birds without breaking down the temporal progression of the first or second 1.5 hours of our 3-hour trials, we found no significant differences in flight—the variable effect of tagging, as a trial progressed, was only elucidated when also considering treatment number. As such, birds flew significantly more in their second treatment, and more when they were untagged (although not significantly so). The importance of this additional covariate was borne out in the second-best and sole competitive model, which included only treatment number. It is possible that, as the birds became more accustomed to the enclosure (reduced neophobia), they grew more comfortable exploring [64]; research has shown that “approach latency” (the speeds at which animals approach novelty) can vary between avian species [65]. Conversely, it is possible that the hummingbirds experienced increasing stress as the trials went on due to additional time in captivity—chronic stress experiments have demonstrated that corticosterone and daytime activity levels can steadily increase to peak in birds after multiple days in captivity [66]—although we did not observe any signs of undue distress (as outlined by Russell et al. [46]) in our video analyses. Furthermore, the acute stress of capturing, handling, and processing in between treatments [67,68] may have increased individuals’ agitation [69], compelling them to remain in a high-alert state throughout their second treatment.

Individual weight, on the other hand, had the highest predictive accuracy of the amount of time that the birds spent feeding, such that increasingly lighter birds fed increasingly more—a significant correlation. It is possible that the heavier hummingbirds we captured had larger energy reserves (e.g., nectar in their crop or digestive system), and thus had less need to replenish their stores [70], whereas the lighter hummingbirds were able to capitalize on an undefended resource through hyperphagia [71,72]. Hummingbirds have an energetic incentive to maximize their short-term nectar intake, particularly when their stores are low [70], allowing them to better accumulate fat [73] and reducing the need for risky torpor [74]. Pre-trial, individuals differed among themselves by up to 24% in bodyweight (7.12–9.31 grams), and over the course of the experiment consumed vastly different quantities of nectar—as little as 3.7% (0.28 grams by a 7.48-gram bird) and as much as 85% (6.62 grams by a 7.77-gram bird) of total bodyweight. Given that it has been speculated that hummingbirds can consume up to three times their bodyweight in a day [75], these results are not necessarily surprising, but rather serve to highlight the extreme level of inter-individual behavioral variation across a sample of adult, male-plumaged hummingbirds from within a single species.

The length of time that individuals spent preening was most accurately predicted through the additive effects of period of day (morning, afternoon, or early evening) and treatment type. This finding is similar in principle to that of flight behavior: if we compared all the time budgets of tagged and untagged birds in a single temporal “bin” (across the full day), we found no significant differences in preening. However, after accounting for the additional covariate of time, the effect of tagging seemingly varied as the day progressed. In this way, preening was significantly greater in tagged individuals, and throughout successively later periods of the day (with increases in preening duration from morning to afternoon, and from afternoon to evening). The reasoning for this relationship is less clear. Reported trends in hummingbird activity as they relate to time of day are somewhat conflicting, with some studies suggesting that feeding activity— which will detract from possible preening time—peaks in morning and evening [76], and others demonstrating steady decreases in feeding activity throughout the day, specifically in the case of *ad libitum* nectar provision [77,78]. Yet there is little information regarding how time spent preening relates to time of day, and further study is warranted. Zenzal et al. [24] point out that birds equipped with auxiliary markers of any kind will, for an unpredictable period, exhibit temporary increases in preening levels before resuming habitual behavior; our results suggest that prolonged monitoring can help to explore circadian effects on time budgets, especially as they relate to biologging ethics.

Based on our inter-rater reliability analysis, our measurements were highly reliable from scorer to scorer, suggesting a well-defined ethogram and little ambiguity in our behavioral categories [57]. The stark contrast between the majority of the birds’ behaviors (e.g., perching versus flying) likely helped to reduce measurement error. With that said, we found the video-analysis stage to be painstaking, owing to the majority of birds undertaking hundreds of rapid flights in each 1-hour treatment (an average of 339 ± 230 when tagged and 362 ± 241 when untagged), as well as due to software-related issues (e.g., BORIS crashing often).

### Aviary Tests

Our research procedure also enabled us to explore the safety of our backpack harness design. As biologger attachment design can play a major role in tag-induced entanglement and mortality [40], we focused on a non-experimental but observational approach to reflect (as closely as possible) routine hummingbird behavior while in an aviary-like setting. Due to the potential for hummingbirds to enter nightly torpor, including tropical species [74], it was also critical to test the potential for overnight device effects. Here, we found that no hummingbirds suffered from vegetative entanglement—even during our recapture efforts at the end of each trial, which we considered a proxy for their ability to evade large predators—nor did we observe any negative effects from overnight wear. In our pilot-version of this study in Ecuador, we similarly found that the three Green-crowned Brilliants that we equipped with identical backpack harnesses for overnight testing suffered no discernible device effects; this species is similarly sized to the Black-throated Mango (our tested brilliants weighed 8.34 ± 0.42 grams).

### Limitations and Future Directions

We currently lack the tools to continuously monitor the behavior of free-living (uncaptured) hummingbirds [79], particularly down to the sub-second level or in ways that will enable us to compare between tagged and untagged individuals (e.g., individuals to act as controls against birds equipped with accelerometers). Thus, rigorous studies of hummingbird biologging ethics must be conducted in confined settings; however, captivity can have significant effects on avian behavior [80,81]. It is possible that our findings, while encouraging, are biased as a result of this limitation [24], and we advise caution when extrapolating them to entirely wild studies. Although our aviary enclosure was more reflective of a hummingbird’s everyday habitat and may have allowed for more “natural” behavior, outdoor filming would have posed both logistical (e.g., camera exposure to rain and humidity [82,83]) and analytical (e.g., visually identifying hummingbirds against non-uniform or low-contrast backgrounds [84,85]) challenges. It is also worth noting that beyond the ordinary stress induced by capture and captivity, the birds’ recent PIT-tagging experience (a surgical procedure) may have exacerbated stress levels [86] and therefore affected behavior. Nonetheless, aside from two individuals PIT-tagged in a prior year, we worked solely with individuals that we had PIT-tagged within the previous five weeks, thereby controlling for this potential effect as much as feasibly possible.

In general, follow-up studies could take a multitude of valuable directions. Incorporating additional aspects of hummingbird biology beyond behavior—such as biomechanics through the use of wind tunnels [6,26], or maneuverability via flight-trajectory tracking within enclosures [87]—would help address other pressing concerns, including escape response speeds and acceleration/turnability differences between tagged and untagged individuals as proxies for survivability [88]. High-speed cameras, meanwhile, would further improve frame-by-frame inter-rater reliability [89], while additionally allowing researchers to conduct straightforward analyses of wingbeat frequency and stroke amplitude (e.g., [21,90]), to examine how hummingbirds might kinematically compensate for the additional weightload of biologgers. Longer-term biologging studies, on the other hand, would allow for a much more comprehensive examination of skin abrasion, which may only be evident after prolonged exposure to a device; attempting to recapture tagged individuals at various intervals after release—days, weeks, months, and even years (such as in [35])—would be extremely informative. Finally, we note that high-quality labelled time budget datasets, such as the one produced through this study (which has a consistent background and focal organism), would provide necessary training and testing data for machine learning algorithms in identifying animal behaviors, opening the door to fresh advancements and collaborations between ecology and computer vision alike [91].

Further studies may also benefit from an updated harness design. While we did not observe any negative harness effects during the course of this experiment, on one occasion with our earlier tests in Ecuador, we observed a White-necked Jacobin attempting to preen under the neck-loop of the harness, at which point its bill became stuck and the bird could not extract it. While White-necked Jacobins have shorter bills than Black-throated Mangoes (our tested birds had exposed culmen lengths of 16.63 ± 0.80 mm *versus* 22.44 ± 1.01 mm, respectively), we would recommend a backpack harness design that does not incorporate a neck-loop. One that conjoins over the sternum to ensure security instead—similar to those used with many raptors [92,93]—may be more suitable; however, a thorough ethical study of this approach is warranted. Regardless of the design, any space between the bird’s body and the harness itself—with or without the neck-loop—may lead to entanglement issues, even if it is as small as a few millimeters (as per the recommendations of Williamson and Witt [35]). Though the birds were able to navigate and avoid the vegetation in our aviary-style enclosure, even during our recapture efforts, our personal observations suggest that hummingbirds are least aware of their surroundings and most likely to come into contact with other objects while in combat with rival individuals. Given that hummingbirds—particularly territorial individuals, such as many of the Black-throated Mangoes [94]—rely on extreme maneuverability, agility, and energy-use for frequent aggressive encounters [36,95], we propose that a valuable follow-up entanglement test would also involve dyadic “competition trials” that spur agonistic behavior, like those designed by Gaffney [96].

In general, this research highlights the need for not only enhanced biologging ethics, but also baseline behavioral information of hummingbirds beyond those that are typically studied in captivity—that is, the North American species hailing from the Bee clade (e.g., [97–101]). Through our work, we have experimentally described the behavior of a species from both a clade (Mangoes, Polytminae) and a region (Colombian Andes) that have garnered much less research. Yet the Black-throated Mango is merely one of Colombia’s roughly 185 species of hummingbirds—the greatest number of any country in the world [102]—and the approximate 366 species ranging across the Americas [103]. We still face a pressing need for basic life history data, and an improved understanding, of these morphologically varied and numerous vertebrate pollinators [36,103–105].

## CONCLUSIONS

Our research approach allowed us to obtain detailed information on hummingbird time budgets; we found no significant differences between the durations, number of bouts, or bout lengths of behaviors exhibited by adult, male-plumaged Black-throated Mangoes when they were and were not equipped with radio-transmitter backpack harnesses. Rather, we found that other variables (whether the birds were undergoing their first or second treatment, the weight of the birds themselves, and the time of day) better predicted the amount of time birds spent performing specific behaviors, either in and of themselves or when considered additively with treatment type (whether the birds were tagged or untagged). Our tests are easy to replicate and/or adjust depending on species-specific needs, providing a remote- and field-accessible approach that can act as a pre-release checkpoint when tagging small-bodied birds—a sorely needed addition to the field of biologging ethics. With that said, we advise caution when considering these results beyond the scope of captive experiments.

## ABBREVIATIONS

-**PIT:** passive integrated transponder
-**RFID:** radio-frequency identification
-**BORIS:** Behavioral Observation Research Interactive Software
-**AICc:** Akaike’s information criterion corrected for small sample sizes

## DECLARATIONS

### Ethics approval and consent to participate

We conducted all work under the approval of the University of Washington’s Institutional Animal Care and Use Committee (IACUC protocol number: 4498-05), and our research permit was issued by Corporación Autónoma Regional de Cundinamarca (permit number: 50227001473). All Ecuadorian pilot research was conducted under the approval and permission of Ministerio del Ambiente y Agua (permit ID: MAAE-ARSFC-2021-1126).

### Consent for publication

Not applicable.

### Availability of data and materials

The datasets and code generated from our study are available in the Dryad repository, [WEB LINK TO DATASETS].

### Authors’ contributions

Lead author AJS led the investigation, project administration, validation, visualization, statistical analysis, figure/table preparation, and writing the original draft of the manuscript; senior author ARG led supervision efforts, and co-led project conceptualization and funding acquisition efforts alongside AJS. AJS, ARG, AMF, and RE established methodology; YT led custom chassis design efforts alongside AJS. Fieldwork was conducted by AJS, AMF, MAM, NTC, and NB. AJS, AMF, and YT led data curation, and all video processing was conducted by AJS, AMF, AEC, SGM, LH, ANC, YS, and OAC. All authors read, provided feedback for, and approved the final manuscript.

## Acknowledgements

We are deeply grateful to Dr. Sarah Converse, Dr. Beth Gardner, Felipe Garzón Agudelo, Deryk Tolman, and Dr. Jay Falk for advice and input on our statistical analyses, as well as late-night ggplot troubleshooting. We also thank Kenna Dailey and Jonathan Bristle for additional video processing assistance, as well as Lucero, Parmenio, and Mary Simbaqueba for unfailingly maintaining feeders and supporting our field team at Centro de Investigación Colibrí Gorriazul. Special thanks to Wilfrido Vaca, Christian Montalvo, and Leonel Montalvo for their support at Un poco del Chocó Reserva y Estación Biológica for the pilot version of this study described in the supplementary material. Last but not least, we are indebted to all hummingbirds that participated in this experiment.

