## Additional File 1 - Supplementary Text, Figures, and Tables for "Tiny radio-tag backpacks impact, but do not significantly affect, hummingbird time budgets in captivity"

### Ecuador Pilot.

#### *Study Site, Species, & Methods.*

In the dry season of August and September 2022, we conducted an observational pilot study of radio-transmitter backpack harness ethics at Un poco del Chocó Estación Biológica (UPDC). At roughly 1,200 m elevation in a transition zone between the Chocó and Tropical Andes hotspots, UPDC is situated alongside the Pachijal River in moderately disturbed, primary lower montane rainforest. We chose to study the Green-crowned Brilliant (*Heliodoxa jacula*), White-necked Jacobin (*Florisuga mellivora*), and White-tipped Sicklebill (*Eutoxeres aquila*) on account of their large body sizes and their varied systematics (each species occupies one of the nine major clades of hummingbirds: Brilliants, Topazes, and Hermits, respectively [1]). In this case, we did not selectively capture adults or males (see **Table S1**). While we sexed individuals based on plumage characteristics, we note that there are recorded cases of male-plumaged females in White-necked Jacobins [2], and that White-tipped Sicklebills are sexually monomorphic [3].

For this experiment, we captured, measured, and radio-transmitter harnessed all hummingbirds as outlined in the main text, but here we identified individuals based on bands as opposed to passive integrated transponders. Our experimental setup was a similar, reduced version of that described in the main text: we constructed our flight arena (Coleman tent) inside the research center, containing a hummingbird feeder (provided sucrose solution measured with a refractometer at the beginning of each trial) and a perch. At the start and end of each trial, we recorded the temperature at the peak of the tent interior using a handheld thermometer. We next released the tagged individual into the flight tent, then manually observed and logged the presence of our behaviors of interest (flying, perching, feeding, and preening) over the trial duration as follows: continuously for hours 0–2, and, if applicable (see **Table S1**), 5-minute checks once every 15 minutes for hours 2–4, 5-minute checks once every 30 minutes for hours 4–6, and then 10-minute checks once every 60 minutes for the remainder of of the trial. In the case of overnight trials, however, we left birds undisturbed until early morning after determining that they had fallen asleep (using a red light to minimize any disturbance).

Note that, as we used a hummingbird feeder as opposed to a syringe, birds perched rather than hovered while feeding. Additionally, given that sicklebills are not known to visit feeders, and the ones in our trial were no exception, we fed each individual by hand once every 30 minutes—an interval that we selected to balance sufficient energetic intake against the stress of recapture inside the tent.

#### *Results & Significance.*

In total, we captured, tagged, and monitored 6 Green-crowned Brilliants (1 adult female, 4 adult males, 1 juvenile of unknown sex), 4 White-necked Jacobins (4 adult males), and 3 White-tipped Sicklebills (all juveniles of unknown sex). Given that this study was observational and our methods differed substantially from those implemented in Colombia, we have elected to reserve our basic results to the supplementary materials here (**Table S1**), and we include them primarily to showcase that there were no drastic behavioral deviations from that which we observed with Black-throated Mangoes, despite the birds' systematic differences.

Average trial lengths, in hours, were as follows: 11.50 ± 10.91 for brilliants, 1.56 ± 0.89 for jacobins, and 4.00 ± 2.00 for sicklebills. While brilliants lost an average of 0.06 ± 0.20 grams across their trials, jacobins and sicklebills gained an average of 0.17 ± 0.15 and 0.74 ± 0.67 grams, respectively. Out of these 13 total individuals, 3 brilliants and 1 jacobin did not preen at all. This result is not inconsistent with what we found in Colombia: 12 of our 25 mangoes exhibited no preening during their tagged observation period (although it is possible that they preened during their 30-minute acclimation period, which we did not film). Beyond this behavior, all individuals exhibited flying, perching, and autonomous feeding (except one, see below), or, in the case of sicklebills, were hand-fed. None of our birds, including those undergoing overnight trials, displayed any of the signs of distress characterized by Russell et al. [4].

As described in the main text, we did observe one instance of harness issues, when a White-necked Jacobin's attempts to preen beneath the harness (sized according to the suggestions of Williamson and Witt [5]) resulted in its bill becoming stuck under the StretchMagic. Indeed, after immediately removing the harness and replacing it with a snugger one, the jacobin began preening again and became stuck once more moments later. We removed this second harness and released the bird promptly, resulting in a curtailed, 0.22-hour trial wherein the bird did not feed at all. Based on these events, we suspect that, owing to birds' many cervical vertebrae [6] and subsequent capacity for head flexion [7], there is an inherent risk to utilizing three-loop harnesses with such proportionally long-billed birds.

Although there have been some studies of Green-crowned Brilliant and White-necked Jacobin behavior, they are largely focused on aggression and dominance [2,8,9]. Research on White-tipped Sicklebills, meanwhile, has been more limited to sighting and feeding records [10–12], given the elusive nature of the sicklebill species [7,13]. While our results do not provide in-depth insight into the daily behaviors of these species either, they are a first step towards understanding how diverse hummingbirds may respond to biologging devices. We strongly believe that comprehensive studies of species-specific effects in relation to biologgers are not only warranted, but critical.

### Harness Design.

#### *Assemblage.*

Adapting from Williamson and Witt [5], we designed our harness to operate with a device that **a)** is solar-powered and can withstand multi-year deployments in the field, and **b)** does not come with prefabricated loops—and thus, any long-term means of attachment to an individual bird. As such, we opted for a top and bottom custom chassis (structural framework) that we could thread around our radio-transmitter (Cellular Tracking Technologies "LifeTag"), effectively pinning it in place while keeping the solar cell fully exposed. We designed our chassis to ensure minimal chafing and drag [14], emphasizing a thin, streamlined nature, rounded corners, and open spaces where possible, to reduce weight without compromising durability.

We designed each chassis (14 mm wide, 22 mm long) individually in Autodesk AutoCAD, a 2D drafting software with DXF file support (other free and open-source options include QCAD and Inkscape), to include **1)** 1-mm diameter holes for the harness and thread, and **2)** each tag's unique identification number (**Fig. S4**). Using a diode-pumped, solid-state, pulsatile laser operating at a wavelength of 355 nm, with a spot size (minimum cut width) of 10 μm, we laser cut each chassis from carbon fiber reinforced polymer (CFRP), which was fabricated with a layup of "prepreg" (fibers pre-impregnated with a thermosetting resin matrix) 27 gsm carbon fiber sheets. We stacked 3 sheets and cured them at 80 psi/150 °C, in 0-90-0 configuration to make the chassis stronger in one direction than the other (more rigid along the length, 0°; more flexible along the width, 90°). The resulting cured sheet is ~90 μm in thickness. It is worth noting that LifeTags are manufactured with a thin coating of epoxy for waterproofing purposes, which acts as an insulating barrier between the conductive CFRP chassis and the tags' circuit components.

We recommend conducting assembly in a setting with access to tools such as a fine-precision scale (~10 mg accuracy), which we used to weigh the tag throughout the construction process. For thread, we chose ultrathin, braided fishing line (KastPro), with a minute application of cyanoacrylate superglue at the tip to avoid fraying. Leaving the four corner holes open for the harness material (0.7 mm StretchMagic), we looped the fishing line through the chassis holes dotted around the LifeTag's perimeter, particularly across its width and in diagonals across the sides, to ensure tight contact between each chassis layer and the tag (**Fig. 1** main text). Throughout this process, we used blunt-end (to avoid fraying), electrostatic-discharge-safe tweezers to guide the fishing line. We opted to add glue via a wire applicator to any knots tied with the fishing line, major points where the thread interacted with the tag, and any places where the tag interacted heavily with the chassis. *Nota bene*: we previously used marine epoxy (UV- and water-resistant) but have since discovered that it is heavy, short-lived, and peels easily.

All told, the addition of the chassis, thread, and glue added a total of ~53 mg to each tag (**Table 1** main text).

#### *Significance and Future Research.*

With the increasing accessibility of miniature animal tracking devices, researchers must continue to refine safe, ultralight attachment methods [5]. When fabricating micromaterials for wildlife applications, there is a continuous tradeoff between weight, durability, and ergonomics (e.g., an overly thin harness thread could cut into a bird's skin, while a slim and narrow chassis, prone to breakage, could impose fewer deleterious device effects). When working with small-bodied birds or insects, there are often few alternatives to glue-on attachment methods, which have unpredictable longevity—although there are countermeasures to increase retention, such as plucking feathers for direct contact with the bird's skin, and gluing or sewing the tag to a gauze or fabric base to increase the connected surface area [15]. In this way, glue-based attachment may be sufficient for miniature radio-transmitters with rapid beep rates and correspondingly short lifespans (< 2–3 months; Lotek, Cellular Tracking Technologies).

In solar-powered tags, however, multi-year deployments are entirely possible (as they are for GPS tags, e.g., [5]), and a flexible, alternative design is essential. As the addition of loops/tubes for harness threading can be too complicated for the manufacturing process or can make tags prohibitively heavy, our chassis design minimizes weight while maximizing longevity and UV capture. CFRPs have a high strength-to-weight ratio—which allows for a very strong yet lightweight chassis—and have a high corrosion resistance, making the chassis resilient to the elements. The combination of cyanoacrylate glue and thread also ensures a longer lifespan, as the former is brittle once cured and does not hold well to bending stresses, while the latter could unravel without additional securement. This ultra-lightweight chassis design, which is easily customizable depending on research needs and tag size/shape, allows us to study a greater expanse of small, previously untestable animals.

Given the rapid technological advancement in the world of biologging [16], we are confident that tags and their associated attachment methods (such as our chassis) can continue to be further improved and streamlined, not only in weight, but also in footprint. Researchers biologging ever-smaller volant animals must ensure that the full range of wing amplitude (e.g., see [17] for hummingbird-specific figures) is in no way obstructed by their devices. Indeed, since conducting this study in 2022, commercially available tags have already been greatly miniaturized, making the tagging process for animals—flighted or not—much safer and more straightforward. As an additional note, although Williamson and Witt [5] found no evidence of deleterious effects for hummingbirds wearing backpack harnesses after 1–2 years, we advise minimizing deployments and maximizing recapture efforts at the termination of each study, owing to the ethical trade-off of individuals wearing tags for multiple years, or even a lifetime.

### Video (caption).

**Additional File 2: Video S1.** Sample behaviors delineated in BORIS video processing. Video clips include flying, hover-feeding, preening, and perching (2 bouts).

### Figures.

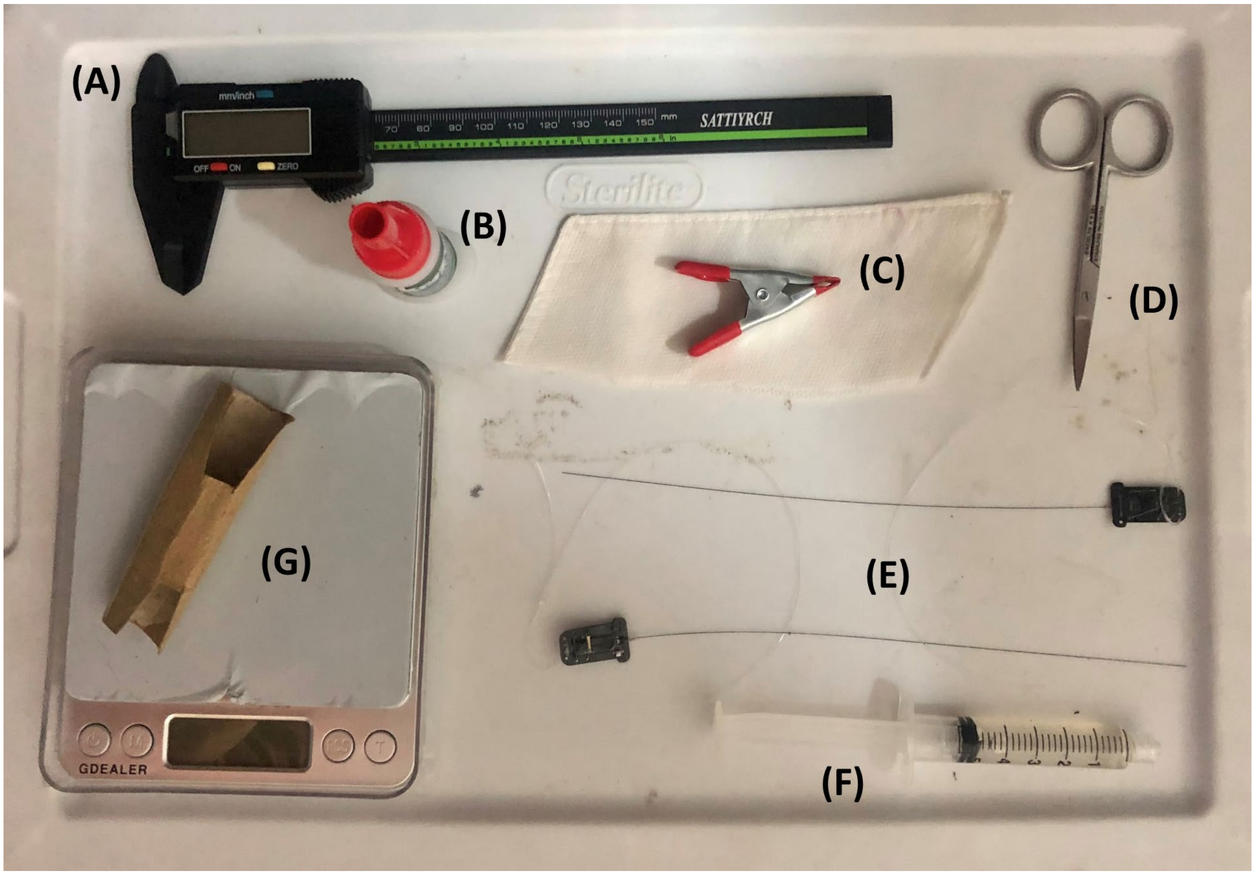


**Figure S1.** Materials used for harness application process. From top-left clockwise: **(A)** calipers to measure harness, **(B)** superglue to secure knots, **(C)** cloth strip and clamp to swaddle bird while tying final harness knot, **(D)** scissors to trim StretchMagic harness once tied, **(E)** LifeTags with prefabricated neck-loops of two sizes to customize for each individual bird, **(F)** prepared syringe with sugar water for if the individual showed any sign of stress (e.g., blinking slowly), and **(G)** scale with cardboard tube to weigh each individual (cutouts were to accommodate the tag on the back post-harnessing).

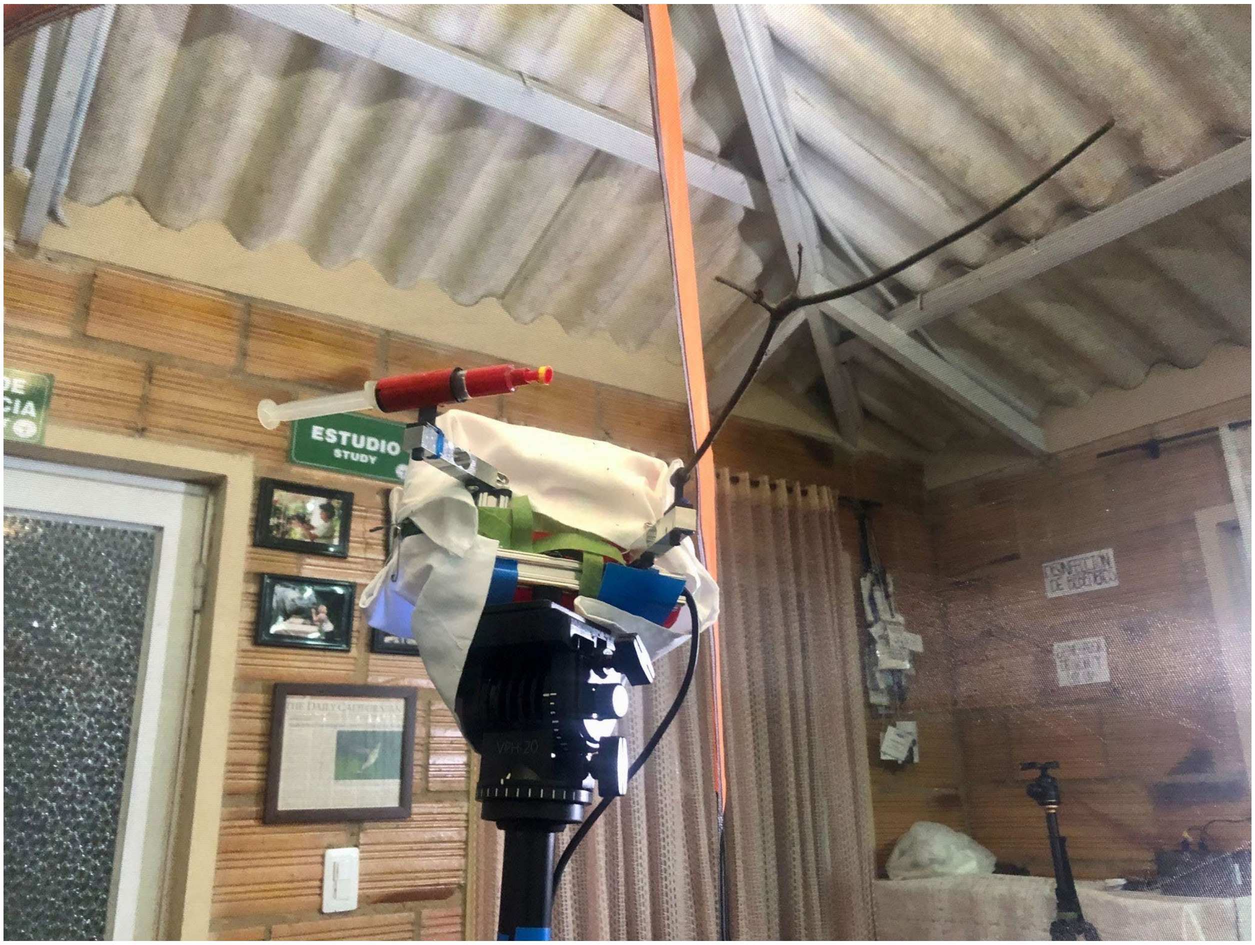


**Figure S2.** Load cell setup in enclosed-environment tent. We coated the syringe on the left-hand side with red tape, adding yellow tape to its tip to mimic floral appearances and draw the hummingbird's attention more readily. We deliberately spaced the perch (right-hand side) far enough from the feeder such that the bird could not perch and feed simultaneously.

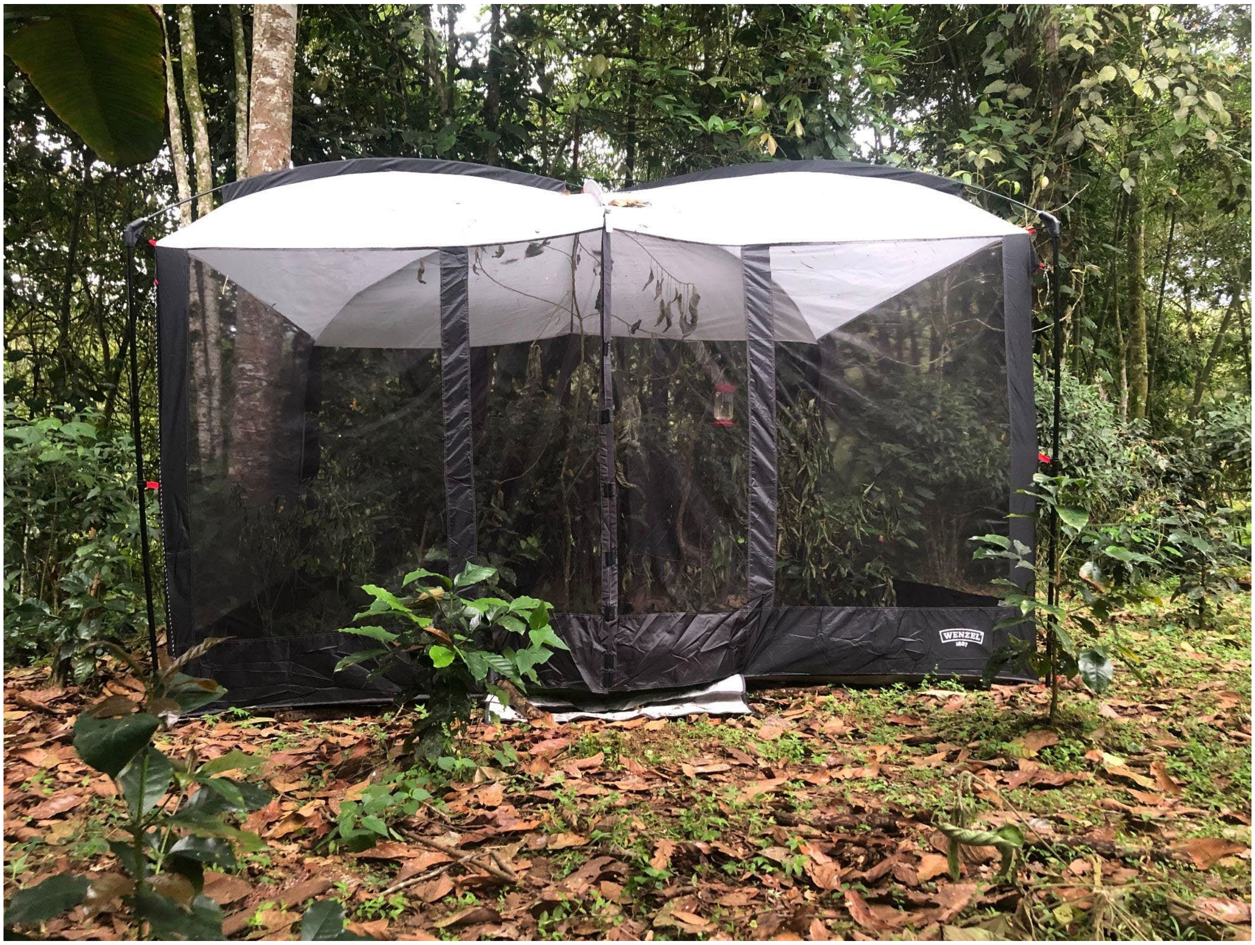

**Figure S3.** Aviary-style enclosure (3.3 x 2.7 x 2.0 m) used for entanglement and overnight harness tests. Feeders and local vegetation are included inside.

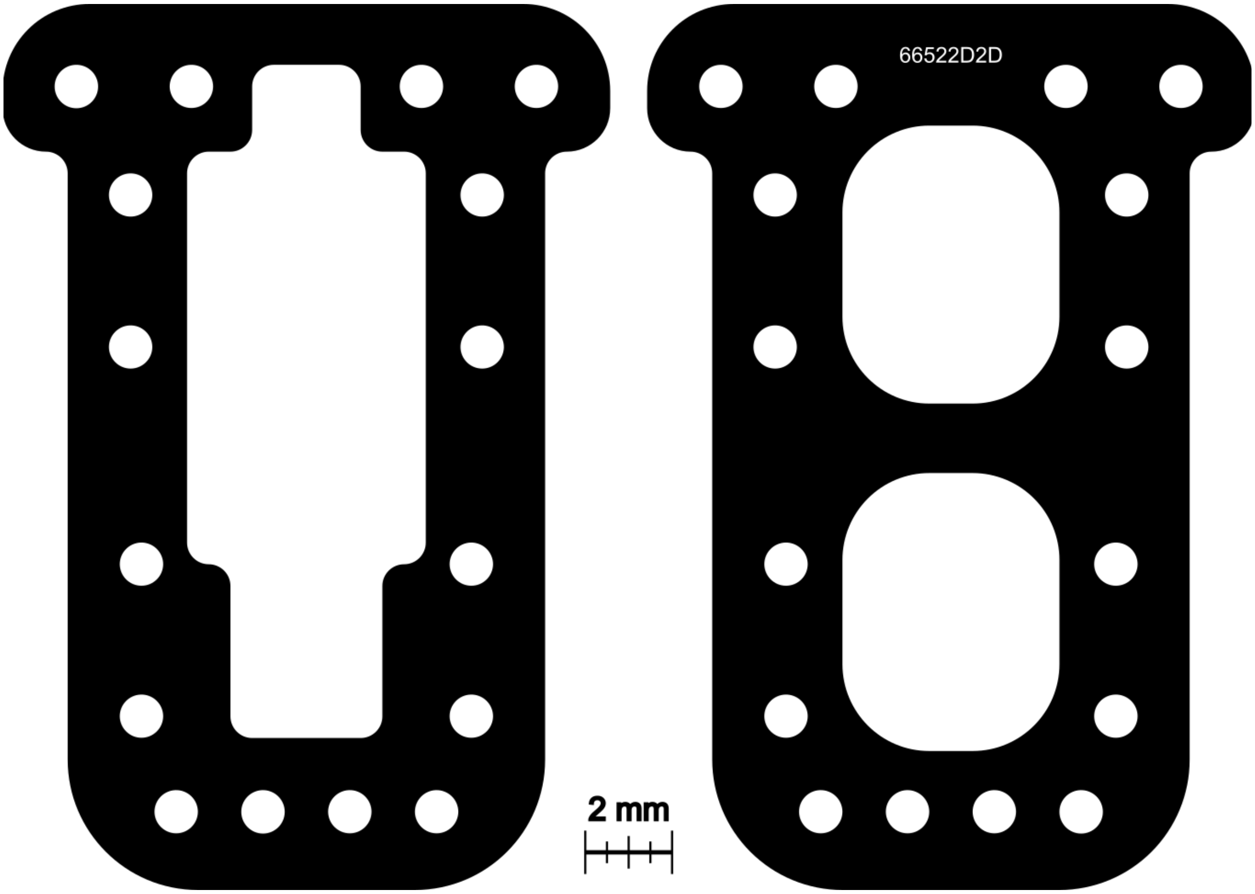


**Figure S4.** Example AutoCAD mock-up for top and bottom chassis of a specific radio-transmitter.

Tables.

**Table S1.** Overview of Ecuadorian pilot study. Column headings represent: species names in Latin (*Species*); bird's sex classification of F. for female, M. for Male, or U. for unknown (*Sex*); bird's age classification of Ad. for adult or Juv. for juvenile (*Age*); trial replicate for a given species (*Trial*); total time in hours that each bird was tested wearing the backpack harness, with * indicating that the trial included overnight testing (*Trial length [h]*); trial's temporal classification of Morn. for morning, Aft. for afternoon, or Eve. for evening, or some combination thereof (*Period of day*); temperature in Celsius within the tent, averaged across beginning and end of each trial (*Temp. [°C]*); w/w sugar-water content of provided nectar (*% sucrose*); behaviors present throughout trial, checked with X to indicate presence, - to indicate absence, or Handfed to indicate the individual was fed by hand (*Behaviors Exhibited: Flying, Perching, Feeding, Preening*); any point that an individual's bill became caught in the neck-loop of the harness (*Harness Issues*), and the amount of weight in grams that the bird gained/lost throughout the trial (*Weight change [g]*).

| Species | Sex | Age | Trial | Trial length (h) | Period of day | Temp. (°C) | % sucrose | Behaviors Exhibited | | | | Harness issues | Weight change (g) |
|---|---|---|---|---|---|---|---|---|---|---|---|---|---|
| | | | | | | | | Flying | Perching | Feeding | Preening | | |
| *H. jacula* | F. | Ad. | 1 | 2 | Aft. | 26.2 | 22 | X | X | X | - | - | + 0.31 |
| | M. | Ad. | 2 | 2 | Aft. | 23.6 | 22 | X | X | X | - | - | - 0.09 |
| | M. | Ad. | 3 | 2 | Morn. | 25.0 | 22 | X | X | X | - | - | - 0.09 |
| | M. | Ad. | 4 | 15* | Eve – Morn. | 18.6 | 22 | X | X | X | X | - | - 0.27 |
| | U. | Juv. | 5 | 24* | All | 18.3 | 22 | X | X | X | X | - | - 0.18 |
| | M. | Ad. | 6 | 24* | All | 19.8 | 22 | X | X | X | X | - | - 0.01 |
| *F. mellivora* | M. | Ad. | 1 | 2 | Morn. | 27.6 | 21 | X | X | X | X | - | + 0.06 |
| | M. | Ad. | 2 | 2 | Eve. | 21.2 | 21 | X | X | X | - | - | + 0.32 |
| | M. | Ad. | 3 | 2 | Morn. | 18.8 | 22 | X | X | X | X | - | + 0.29 |
| | M. | Ad. | 4 | 0.22 | Aft. | 23.8 | 22 | X | X | - | X | X | - 0.02 |
| *E. aquila* | U. | Juv. | 1 | 2 | Aft. | 24.0 | 21 | X | X | Handfed | X | - | - 0.09 |
| | U. | Juv. | 2 | 4 | Aft. | 22.0 | 21 | X | X | Handfed | X | - | + 1.43 |
| | U. | Juv. | 3 | 6 | Morn. – Aft. | 21.6 | 23 | X | X | Handfed | X | - | + 0.71 |

**Table S2.** Model rankings for different behavioral durations (flight, hover-feeding, and preening), with each single fixed-effect model including the random effect of individual. Error model structures and zero-inflation terms are specified. Column headings represent: model rank (*Rank*), fixed effect included in model (*Model*) number of estimated parameters (*k*), log-likelihood (*LL*), Akaike's information criterion corrected for small sample sizes (*AICc*), delta AICc score (*ΔAICc*), and Akaike weight (*Weight*; indicates the level of support in favor of any given model being the most parsimonious candidate). The rankings of all models that had a significant likelihood ratio test (performed against the null) are bolded.

| Rank | Model | k | LL | AICc | ΔAICc | Weight |
|---|---|---|---|---|---|---|
| | **Flight (Gaussian)** | | | | | |
| **1** | Treatment Number + Treatment Type | 5 | -380.88 | 773.13 | 0.00 | 0.45 |
| **2** | Treatment Number | 4 | -382.65 | 774.20 | 1.07 | 0.26 |
| **3** | Treatment Number * Treatment Type | 6 | -380.87 | 775.68 | 2.55 | 0.12 |
| 4 | Treatment Type | 4 | -384.41 | 777.71 | 4.58 | 0.05 |
| 5 | Null (random effect only) | 3 | -385.62 | 777.77 | 4.64 | 0.04 |
| 6 | % Sucrose | 4 | -384.76 | 778.40 | 5.27 | 0.03 |
| 7 | Bird Weight | 4 | -385.00 | 778.89 | 5.76 | 0.03 |
| 8 | Temperature | 4 | -385.53 | 779.95 | 6.82 | 0.01 |
| 9 | Time of Day | 5 | -385.08 | 781.53 | 8.40 | 0.01 |
| | **Feeding (Tweedie, zero-inflated)** | | | | | |
| **1** | Bird Weight | 6 | -200.26 | 414.47 | 0.00 | 0.95 |
| 2 | Null (random effect only) | 5 | -205.36 | 422.08 | 7.62 | 0.02 |
| 3 | Treatment Number + Bird Weight | 7 | -202.99 | 422.65 | 8.18 | 0.02 |
| 4 | Treatment Number * Bird Weight | 8 | -202.98 | 425.48 | 11.01 | 0.00 |
| 5 | Treatment Type | 6 | -205.81 | 425.57 | 11.10 | 0.00 |
| 6 | % Sucrose | 6 | -206.08 | 426.12 | 11.65 | 0.00 |
| 7 | Time of Day | 7 | -205.56 | 427.79 | 13.33 | 0.00 |
| 8 | Temperature | 6 | -207.01 | 427.97 | 13.51 | 0.00 |
| 9 | Treatment Number | 6 | -207.17 | 428.30 | 13.84 | 0.00 |
| | **Preening (Tweedie, zero-inflated)** | | | | | |
| **1** | Treatment Type + Time of Day | 8 | -168.03 | 355.57 | 0.00 | 0.90 |
| **2** | Treatment Type * Time of Day | 10 | -167.56 | 360.76 | 5.19 | 0.07 |
| **3** | Time of Day | 7 | -173.03 | 362.72 | 7.15 | 0.03 |
| **4** | Treatment Type | 6 | -176.52 | 366.99 | 11.42 | 0.00 |
| 5 | Null (random effect only) | 5 | -179.83 | 371.02 | 15.45 | 0.00 |
| 6 | Treatment Number | 6 | -179.24 | 372.43 | 16.86 | 0.00 |
| 7 | Temperature | 6 | -179.76 | 373.48 | 17.91 | 0.00 |
| 8 | Bird Weight | 6 | -179.80 | 373.55 | 17.98 | 0.00 |
| 9 | % Sucrose | 6 | -179.81 | 373.58 | 18.01 | 0.00 |

**Table S3.** Parameter estimates, standard errors (SE), 95% confidence intervals (CI), and p-values (p) for the top-ranked models for flight, hover-feeding, and preening durations, as determined using Akaike's information criterion adjusted for small sample sizes (AICc).

| Parameter | Estimate | SE | Lower 95% CI | Upper 95% CI | p |
|---|---|---|---|---|---|
| **Flight ~ Treatment Number + Treatment Type (Gaussian)** | | | | | |
| β Intercept | 776.60 | 138.10 | 498.01 | 1055.19 | 2.61e-06 |
| β Untagged | 149.39 | 76.61 | -6.72 | 305.49 | 0.0625 |
| β Treatment 2 | 218.74 | 76.61 | 62.64 | 374.84 | 0.0085 |
| **Feeding ~ Bird Weight (Tweedie, zero-inflated)** | | | | | |
| β Intercept | 5.98 | 0.88 | 4.27 | 7.70 | 8.25e-12 |
| β Weight | -0.29 | 0.11 | -0.51 | -0.07 | 0.0086 |
| **Preening ~ Time of Day + Treatment Type (Tweedie, zero-inflated)** | | | | | |
| β Intercept | 4.74 | 0.36 | 4.04 | 5.44 | < 2e-16 |
| β Untagged | -1.54 | 0.42 | -2.36 | -0.72 | 0.0002 |
| β Morning | -1.84 | 0.56 | -2.94 | -0.74 | 0.0010 |
| β Evening | 0.82 | 0.46 | -0.08 | 1.72 | 0.0746 |

### References.

7. Boehm MMA, Guevara-Apaza D, Jankowski JE, Cronk QCB. Floral phenology of an Andean bellflower and pollination by buff-tailed sicklebill hummingbird. Ecol Evol. 2022;12:e8988.
8. Falk JJ, Rubenstein DR, Rico-Guevara A, Webster MS. Intersexual social dominance mimicry drives female hummingbird polymorphism. Proc R Soc B Biol Sci. 2022;289:20220332.
9. Fernandez-Duque F, Miller ET, Fernandez-Duque M, Falk J, Venable G, Rabinowicz S, et al. Phenotype predicts interspecific dominance hierarchies in a cloud-forest hummingbird guild. Ethology. 2024;130:e13410.
10. Stiles FG. Ecology, Flowering Phenology, and Hummingbird Pollination of Some Costa Rican Heliconia Species. Ecology. 1975;56:285–301.
11. Vigle GO. A Nest of Eutoxeres aquila heterura in Western Ecuador. The Auk. 1982;99:172–3.
12. Rengifo C, Bakermans MH, Puente R, Vitz A, Rodewald AD, Zambrano M. First Record of the White-tipped Sicklebill (Eutoxeres aquila aquila: Trochilidae) for Venezuela. Wilson J Ornithol. 2007;119:292–5.
13. Stiles FG. The Annual Cycle in a Tropical Wet Forest Hummingbird Community. Ibis. 1980;122:322–43.
14. Vandenabeele SP, Shepard ELC, Grémillet D, Butler PJ, Martin GR, Wilson RP. Are bio-telemetric devices a drag? Effects of external tags on the diving behaviour of great cormorants. Mar Ecol Prog Ser. 2015;519:239–49.
15. Hadley AS, Betts MG. Tropical deforestation alters hummingbird movement patterns. Biol Lett. 2009;5:207–10.
16. Kays R, Crofoot MC, Jetz W, Wikelski M. Terrestrial animal tracking as an eye on life and planet. Science. 2015;348:aaa2478.
17. Díaz-Salazar AF, Garzón-Agudelo F, Smiley A, Cadena CD, Rico-Guevara A. Winging it: Hummingbirds alter flying kinematics during molt. Biol Open. 2024;bio.060370.
